# Microglial phagocytosis dysfunction is related to local neuronal activity in a genetic model of epilepsy

**DOI:** 10.1101/2020.05.06.075903

**Authors:** Virginia Sierra-Torre, Ainhoa Plaza-Zabala, Paolo Bonifazi, Oihane Abiega, Irune Díaz-Aparicio, Saara Tegelberg, Anna-Elina Lehesjoki, Jorge Valero, Amanda Sierra

## Abstract

Microglial phagocytosis of apoptotic cells is an essential component of the brain regenerative response in neurodegenerative diseases. Phagocytosis is very efficient in physiological conditions, as well as during apoptotic challenge induced by excitotoxicity or inflammation, but is impaired in mouse and human mesial temporal lobe epilepsy (MTLE). Here we extend our studies to a genetic model of progressive myoclonus epilepsy type 1 (EPM1) in mice lacking cystatin B (CSTB), an inhibitor of cysteine proteases involved in lysosomal proteolysis. We first demonstrated that microglial phagocytosis was impaired in the hippocampus in *Cstb* knock-out (KO) mice when seizures arise and hippocampal atrophy begins, at 1 month of age. To test if this blockage was related to the lack of *Cstb* in microglia, we used an *in vitro* model of phagocytosis and siRNAs to acutely reduce *Cstb* expression but we found no significant effect in the phagocytosis of apoptotic cells. We then tested whether seizures were involved in the phagocytosis impairment, similar to MTLE, and analyzed *Cstb* KO mice before seizures begin, at postnatal day 14. Here, phagocytosis impairment was restricted to the granule neuron layer but not to the subgranular zone, where there are no active neurons. Furthermore, we observed apoptotic cells (both phagocytosed and not phagocytosed) in *Cstb* deficient mice at close proximity to active, cFos^+^ neurons and used mathematical modeling to demonstrate that the physical relationship between apoptotic cells and cFos+ neurons was specific for *Cstb* KO mice. These results suggest a complex crosstalk between apoptosis, phagocytosis and neuronal activity, hinting that local neuronal activity could be related to phagocytosis dysfunction in *Cstb* KO mice. Overall, this data suggest that phagocytosis impairment is an early feature of hippocampal damage in epilepsy and opens novel therapeutic approaches for epileptic patients based on targeting microglial phagocytosis.

## INTRODUCTION

Progressive myoclonus epilepsy 1 (EPM1) or Unverricht-Lundborg disease is a genetic neurodegenerative disease characterized by disabling myoclonus, epileptic seizures and ataxia. The disease is caused by biallelic loss-of-function mutations in the cystatin B gene *(CSTB)* encoding the cystatin B protein^1–3^, a protease inhibitor that regulates and limits the activity of nuclear and cytoplasmic cysteine proteases known as cathepsins^4,5^. *Cstb* knock-out (KO) mice recapitulate key clinical features of EPM1^5,6^, as they develop seizures at one month and progressive ataxia around six months of age^7–10^, consistent with findings in humans. In addition, *Cstb* KO mice present early alterations in the inflammatory response of microglia, the brain immune cells^11–13^. In addition to controlling the release of inflammatory mediators, microglia are also the brain professional macrophages^14^. We have recently reported that in both mouse and human mesial temporal lobe epilepsy (MTLE), microglial phagocytosis of apoptotic cells is impaired^15^, leading us to question whether the phagocytosis impairment would also occur in *Cstb* KO mice.

Phagocytosis is a highly regulated response that prevents the spillover of cytotoxic content that results from the cell death, is immunomodulatory, and actively participates in maintaining tissue homeostasis^14,16,17^. In physiological conditions, microglia are very efficient phagocytes, as their processes constantly scan the brain parenchyma and express a plethora of receptors for “find-me” and “eat-me” signals produced by apoptotic cells^18–21^. Microglia are similarly efficient facing increased apoptotic cell numbers generated during excitotoxicity and inflammation, as they use several strategies to cope with increased apoptosis: 1) recruitment of more phagocytic cells, 2) increase the number of apoptotic cells phagocytosed per microglial cell, and/or 3) increase the microglial numbers^15,22^. As a consequence, microglial phagocytosis is proportional to the number of apoptotic cells, i.e. microglial phagocytosis is tightly coupled to apoptosis^15^. However, during MTLE the hyperactivity of the neuronal network and the massive release of the “find-me” signal ATP during seizures interfered with the recognition by microglia, resulting in delayed dead cell clearance and inflammation^15^.

To test whether microglial phagocytosis was affected in other forms of epilepsy, such as EPM1, we analyzed microglial phagocytosis efficiency in *Cstb* KO mice. Most of the CSTB deficiency -related pathological changes have been reported in the cerebellum and the cortex^7,9–11^ but here we focused on the dentate gyrus (DG) of the hippocampus, in whose subgranular zone (SGZ) there is ongoing production of newborn neurons through adulthood^23^. Most of these newborn neurons naturally undergo apoptosis and are immediately phagocytosed by microglia, allowing us to establish the baseline phagocytosis efficiency^22,24^. We here show hippocampal atrophy and microglial phagocytosis impairment at the age when seizures arise, at postnatal day 30 (P30). We then studied whether this effect was related to a cell-autonomous effect on microglia, using an *in vitro* model of phagocytosis and siRNA-mediated depletion; or was related to altered neuronal function, by studying presymptomatic mice at P14, before seizures arise. We found that in the neurogenic region, the SGZ, phagocytosis was spared; in contrast, phagocytosis was specifically impaired in the granule neuron cell layer. We finally used direct analysis and mathematical modeling to determine that apoptotic cells were in fact in close proximity to cFos^+^, activated neurons, in the absence of seizures. Our data suggests a complex environmental signaling in *Cstb* KO mice that results in granule cell layer-specific microglial phagocytosis impairment and point towards microglia as a potential target of novel treatments aimed at reducing neuronal damage in epilepsy patients.

## METHODS

### Mice

Tissues of *Cstb* KO mice^7^ (129S2/SvHsd5-*Cstb*^tm1Rm^; Jackson Laboratory stock no. #003486) were obtained from the Lehesjoki’s lab at Folkhälsan Research Center and University of Helsinki. Wild-type litter mates with the same genetic background were used as controls. For the FACS experiments, fms-EGFP mice (B6.Cg-Tg(Csf1r-EGFP)1Hume/J; Jackson Laboratory stock #018549), where microglia constitutively express the GFP protein^25,26^ were used. All procedures followed the European Directive 2010/63/EU and NIH guidelines, and were approved by the Animal Ethics Committee of the State Provincial Office of Southern Finland (decisions ESAVI/7039/04.10.03/2012 and ESAVI/10765/04.10.07/2015) and Ethics Committees of the University of the Basque Country EHU/ UPV (Leioa, Spain; CEBA/205/2011, CEBA/206/2011, CEIAB/82/2011, CEIAB/105/2012).

### BV2 cell line

BV2 cells (Interlab Cell Line Collection San Martino-Instituto Scientifico Tumori-Instituto Nazionale per la Ricerca sul Cancro), a cell line derived from raf/myc-immortalized rat neonatal microglia was used to perform the *in vitro* knock-down model of *Cstb.* BV2 cells were grown as an adherent culture in non-coated 90mm^2^ Petri dishes in the presence of 10 mL of culture medium. The culture medium consisted in high glucose Dulbecco’s Modified Eagle’s Medium (DMEM, Invitrogen) supplemented with 10% fetal bovine serum (FBS, GE Healthcare Hyclone) and 1% antibiotic-antimycotic (Gibco). When confluence was reached, cells were trypsinized (Trypsin-EDTA 0,5% no phenol red, Gibco) and replated at 1:5.

### Red Fluorescent SH-SY5Y Cell Line

(Vampire SH-SY5Y). The Vampire SH-SY5Y cell line was developed as a stable transfection of the SH-SY5Y cell line with the red fluorophore tFP602 (InnoProt, P20303). Cells were grown as an adherent culture in non-coated T-75 flasks in the presence of 10 mL of culture medium. The culture medium consisted in high glucose DMEM supplemented with 10% fetal bovine serum (FBS, GE Healthcare Hyclone) 1% antibiotic-antimycotic (Gibco) and 0.25mg/ml Geneticin (G418, Gibco) to select the transfected cells. When confluence was reached, cells were trypsinized (Trypsin-EDTA 0,5% no phenol red, Gibco) and replated at 1:3.

### *In vitro* phagocytosis assay

The protocol for *in vitro* phagocytosis was adapted from^27^. Briefly, BV2 cells were transfected and phagocytosis was performed after 24 hours of expression. Phagocytosis was performed in high glucose DMEM, supplemented 1% antibiotic-antimycotic (Gibco) and 10% fetal bovine serum (FBS, GE Healthcare Hyclone) to ensure the presence of the complement molecules, such as C1q, that have been related to phagocytosis *in vivo^28^* and determine the immunomodulatory outcome of phagocytosis^29^. BV2 cells, transfected with either *Cstb* or scrambled siRNA, were fed for 1 and 4h with Vampire SH-SY5Y, previously treated with staurosporine (STP; 3μM, 4 h; Sigma-Aldrich) to induce apoptosis. The floating fraction of the dead cells was collected and added to the BV2 cells in a 1:1 proportion after their visualization and quantification with trypan blue in a Neubauer chamber.

### Immunofluorescence

Six series of 50μm-thick sagittal sections per brain were sectioned using a Leica VT 1200S vibrating blade microtome (Leica Microsystem GmbH, Wetzlar, Germany). Immunostaining was performed following standard procedures^22,27^. Free floating vibratome sections were blocked in permeabilization solution (0.3% Triton X-100, 0.5% BSA in PBS; all from Sigma-Aldrich) for 2 h at room temperature (RT) with gentle shaking, and incubated overnight with the primary antibodies diluted in the permeabilization solution at 4°C. After primary antibody incubation, sections were intensively washed with 0.3% Triton X-100 in PBS and then incubated with secondary antibodies and DAPI (5mg/mL, Sigma-Aldrich) diluted in permeabilization solution for 2h at RT. After washing with PBS, the sections were mounted on glass slides with Dako Cytomation Fluorescent Mounting Medium.

BV2 cells were fixed for 10 min at RT in 4% PFA and then transferred to PBS. Fluorescent immunostaining was performed following standard procedures^27,15^. Coverslips with BV2 cells were blocked in 0.2% Triton X-100, 0.5% BSA in PBS for 1h at RT. The cells were then incubated overnight with primary antibodies in permeabilization solution (0.2% Triton X-100, 0.5% BSA in PBS). After overnight incubation, cells were rinsed in PBS and incubated in the secondary antibodies containing DAPI (5 mg/ml) in the permeabilization solution for 1 h at RT. After washing with PBS, primary cultures were mounted on glass slides with Dako Cytomation Fluorescent Mounting Medium.

### Antibodies

The following antibodies were used; chicken anti-GFP (1:1000, Aves Laboratoies), rabbit anti-Iba1 (1:1000, Wako), goat anti-Iba1 (1:1000, Abcam), rat anti-CD11b (1:200, BioRad), rabbit anti-Ki67 (1:1000, Vector Laboratories), rabbit anti-GFAP (1:1000, Abcam), rabbit anti-cFos (1:750, Santa Cruz Biotechnologies), mouse anti-NeuN (1:750, Millipore). Secondary antibodies coupled to AlexaFluor 488, Rodhamine Red X or AlexaFluor 647 were purchased from Jackson Laboratories.

### Image analysis

All fluorescence immunostaining images were collected using a Leica SP8 laser-scanning microscope using a 40x oil-immersion objective and a z-step of 0.7μm. All images were imported into the Fiji distribution of ImageJ^30^ in Tiff format. Brightness, contrast, and background were adjusted equally for the entire image using the “brightness and contrast” and “levels” controls from the “image/adjustment” set of options without any further modification. For tissue sections, two to three 20μm-thick z-stacks of each section containing the septal hippocampus from one vibratome series were analyzed. For BV2 cell cultures, over 4 −5 z-stacks were obtained per coverslip.

### Phagocytosis analysis *in vivo* and *in vitro*

Apoptotic cells were defined based on their nuclear morphology after DAPI staining as cells in which the chromatin structure (euchromatin and heterochromatin) was lost and appeared condensed and/or fragmented (pyknosis/ karyorrhexis). Phagocytosis was defined as the formation of an enclosed, three-dimensional pouch of microglial processes surrounding an apoptotic cell^22^. In tissue sections, the number of apoptotic cells, phagocytosed cells and microglia were estimated using unbiased stereology in the volume of the DG contained in the z-stack (determined by multiplying the thickness of the stack by the area of the DG at the center of the stack using ImageJ, Fiji). To obtain the total numbers, this density value was then multiplied by the volume of the septal hippocampus (spanning from −1 to −2.5 mm in the AP axes, from bregma; 6 slices in each of the 6 series), which was calculated using Fiji from Zeiss Axiovert epifluorescent microscope images collected at 20X. *In vitro,* the percentage of phagocytic microglia was defined as cells with pouches containing apoptotic Vampire-SH-SY5Y nuclei and/or RFP particles^27^. To calculate the phagocytosis parameters, the following formulas are used^27^:

#### Phagocytic Index

proportion of apoptotic cells engulfed by microglia. Phagocytosed apoptotic cells (apo^ph^); total apoptotic cells (apo^tot^).

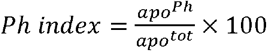

#### Phagocytic capacity

proportion of microglia with one or more phagocytic pouches, each containing one apoptotic cell^22^. Microglia (nig); microglia with one or more phagocytic pouches (Phn).

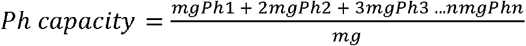

#### Phagocytosis/Apoptosis coupling

net phagocytosis (number of microglia multiplied by their phagocytic capacity) divided by the number of apoptotic cells^15^.

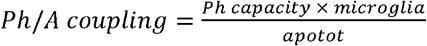

### BV2 cell line transfection

siRNA targeting *Cstb* (*Cstb* siRNA) and scrambled siRNA (shuffled *Cstb* sequence) (Thermo Fisher) were used to transfect BV2 cells. Both *Cstb* and scrambled siRNA were labeled for the immunofluorescence assays using the Silencer Labeling Kit with FAM dye (Invitrogen, AM1634) following the manufacturer’s instructions. BV2 cells were seeded in 24 or 6 well plates 24 hours prior to the transfection assay in high glucose DMEM supplemented with 10% fetal bovine serum (FBS, GE Healthcare Hyclone) and 1% antibiotic-antimycotic (Gibco) to ensure ~80% confluence the day of the experiment. In 24-well plates 30.000 cells per well were seeded and in 6-well plates 150.000 cells per well were seeded. Transfection assay was performed following manufacturer’s instructions. Lipofectamine 2000 (2μg/mL, Invitrogen) and 5nM siRNAs were separately pre-incubated with Opti-MEM (Gibco). Lipofectamine 2000 and siRNAs were left to complex for 20 minutes at room temperature and added to the plated BV2 cells in low glucose DMEM supplemented with 5% fetal bovine serum (FBS, GE Healthcare Hyclone) and no antibiotics to ensure full transfection efficiency. Lipofectamine 2000-siRNAs complexes were added to BV2 cells for 15 hours and then replaced with the regular growth medium, high glucose DMEM supplemented with 10% fetal bovine serum (FBS, GE Healthcare Hyclone) and 1% antibiotic-antimycotic (Gibco). siRNA knock-down was assessed 6, 24 and 48 hours after transfection both by FAM-immunofluorescence and RT-qPCR.

### FACS sorting

Microglia were isolated from 1 month old mouse hippocampi^26,15^. In brief, the brains from fms-EGFP mice were enzymatically disaggregated (enzymatic solution, in mM: 116 NaCl, 5.4 KCl, 26 NaHCO3, 1 NaH2PO4, 1.5 CaCl2, 1 MgSO4, 0.5 EDTA, 25 glucose, 1 L-cysteine; with papain (20 U/ml) and DNase I (150 ul; Invitrogen) for at 37°C for 15 min. The enzymatic digestion was accompanied by mechanical disaggregation through careful pipetting. After homogenization, the cell suspension was filtered through a 40μm nylon strainer to a 50 ml Falcon tube quenched by 5 ml of 20% FBS in HBSS. To further enrich the microglial population, myelin was removed by using Percoll gradients; cells were centrifuged at 200g for 5 min and resuspended in a 20% solution of isotonic Percoll (SIP; 20% in HBSS), obtained from a previous SIP stock (9 parts Percoll per 1 part PBS 10X). Each sample was layered with HBSS poured very slowly by fire-polished pipettes. Afterwards, gradients were centrifuged for 20 min at 200g’s with minimum acceleration and no brake so the interphase was not disrupted. Then, the interphase was removed, cells were washed in HBSS by centrifuging at 200g’s for 5 min and the pellet was resuspended in 500μl of sorting buffer (25 mM HEPES, 5 mM EDTA, 1% BSA, in HBSS). Microglia cell sorting was performed by FACS Jazz (BD Biosciences, in which the population of green fluorescent cells was selected, collected in Lysis Buffer (Qiagen) containing 0.7% -mercaptoethanol and stored at −80°C until processing.

### RNA isolation and retrotranscription

RNA from FACS-sorted microglia was isolated by RNeasy Plus micro kit (Qiagen) according to the manufacturer’s instructions and retrotranscribed using the iScript Advanced cDNA Synthesis Kit (BioRad). RNA from transfected BV2 cells was isolated by RNeasy Plus mini kit (Qiagen) according to the manufacturer’s instructions and retrotranscribed using the Superscript III Reverse Transcriptase (Invitrogen).

### Real-time qPCR

Real-time qPCR was performed following MIQE guidelines (Minimal Information for Publication of Quantitative Real Time Experiments)^31^. Three replicas of 1.5μl of a 1:3 dilution of cDNA were amplified using SsoFast EvaGreen Supermix (BioRad) for FACS-sorted microglia and Power SYBR Green (Bio-Rad) for BV2 transfected cells in a CFX96 Touch Real-Time PCR Detection System (Bio-Rad). The amplification protocol for both enzymes was 3 min 95°C, and 40 cycles of 10 s at 95°C, 30 s at 60°C.

### Primers

Cstb, cathepsins B L and *S* commercial primers were purchased from Sigma-Aldrich (**Table 1**). Their specificity was assessed using melting curves and electrophoresis in 2% agarose gels. For each set of primers, the amplification efficiency was calculated using the software LinRegPCR^32^ or standard curve of 1:2 consecutive dilutions, and was used to calculate the relative amount using the ΔΔCt following formula:

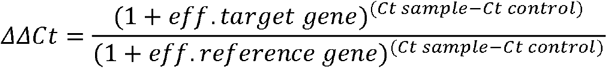

**Table 1.**
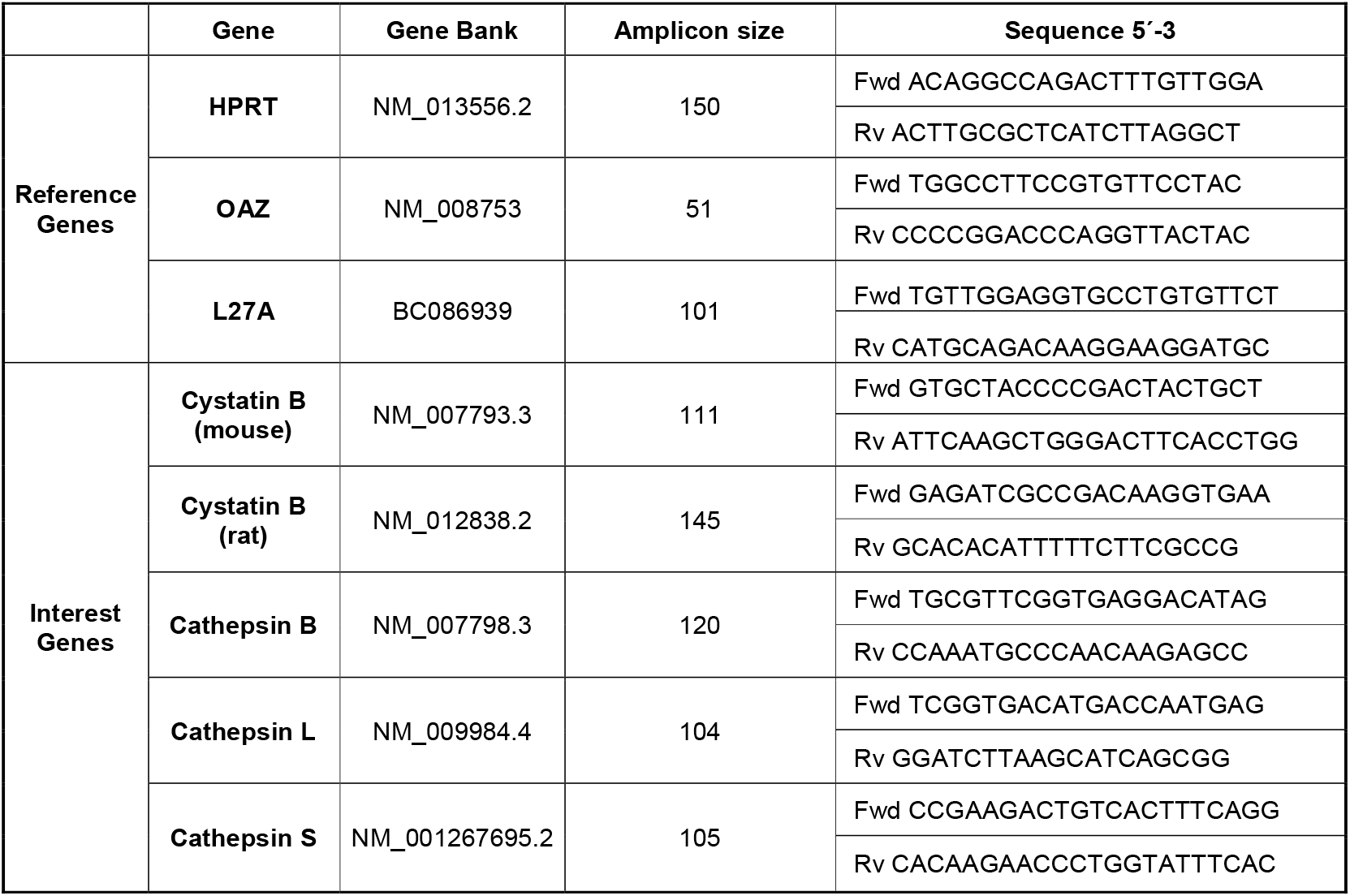
Gene accession number, primer sequence, amplicon size

Up to three independent reference genes were compared: L27A, which encodes a ribosomal protein of the 60S subunit^26^; OAZ-1, which encodes ornithine decarboxylase antizyme, a rate-limiting enzyme in the biosynthesis of polyamines and recently validated as a reference gene in rat and human^33^; and HPRT, which encodes hypoxanthine guanine phosphoribosyl transferase^34^. The expression of L27A, OAZ-1, and HPRT remained constant independently of time and treatments, validating their use as reference genes. The reference gene that rendered lower intragroup variability was used for statistical analysis.

### Direct determination of apoptotic cell to cFos^+^ neurons distances

For each image, the 3D coordinates of cFos^+^ and apoptotic cells of the granule layer (GL) were visually identified and saved using the Point tool of the Fiji distribution of ImageJ. Then, we used these coordinates to estimate the cartesian distances of each apoptotic cell to its closest cFos^+^ neuron (nearest neighbor, NN) and to the nearest border of the image stack (X, Y and Z borders). This direct analysis, however, was hindered by the fact that most apoptotic cells were closer to a border of the z-stack (the upper or bottom Z border in most cases) than they were to a cFos^+^ neuron (**Supplementary Figure 1**). Therefore, the inclusion criteria for analyzing apoptotic-cFos+ distances was: d_cFosNN_<d_border_ (i.e., the calculated distance to cFos NN must be shorter than the distance to the closest border), and apoptotic-cFos+ distances that were further away than the distance to the closest border, which was the case for the vast majority of apoptotic cells, were discarded. We analyzed a total of 25 and 253 apoptotic cells collected from 2 series of sections from the GL of wild-type (WT) and KO mice, respectively. Out of these, only 5 phagocytosed and 15 non phagocytosed cells from n=9 KO mice, and one cell of each type in n= 9 WT mice passed the inclusion criteria. To ensure that our z-stacks did not have an intrinsic bias in either of the cell types, we confirmed that both phagocytosed and non-phagocytosed apoptotic cells were located in z-stacks of similar thickness (23.4 ± 0.6 vs 24.3 ± 0.4 μm, respectively), and therefore had similar chance of detection.

### Modelling apoptotic cell to cFos^+^ neurons distances

To generate the model, we first generated virtual 3D (v3D) cell location grids based on real granule neurons. On these grids, we placed apoptotic cells and cFos^+^ neurons in 10,000 simulations and calculated cumulative probability distribution function of the distance between apoptotic and NN cFos^+^ cells

#### 1. Generation of v3D cell location grids from real WT and KO z-stack images

To model the distances between apoptotic and cFos+ cells we generated v3D grid locations of all granule neurons (NeuN^+^) and apoptotic cells from 16 representative confocal z-stack images from WT and KO mice (8 z-stacks each). For each set of z-stack images, the 3D coordinates of NeuN^+^ granule neurons and apoptotic cells of the GCL were visually identified and saved using the Point tool of Fiji. These coordinates were pooled to generate a v3D grid for each z-stack, on which cFos^+^ and apoptotic cells were later positioned in four different random models (see below). The computations for the models were performed in Matlab (MathWorks, Natick, MA).

#### 2. Number of cFos-positive and apoptotic cells located in each virtual 3D grid

The number of apoptotic cells located in each v3D grid was derived randomly from a Gaussian probability distribution with mean and standard deviation (SD) obtained from the experimental apoptotic cell density (**Table 2**), multiplied by the total number of cells in the grid and finally approximated to the closest integer positive number. In the case of models 1-3, WT- and KO-derived v3D grids were filled according to the corresponding WT and KO densities, respectively. In model 4 we used inverted densities, and filled the WT v3D grid from the KO density, and vice versa for the KO v3D grid. The number of cFos^+^ cells located on the v3D grids either corresponded to the real count in the correspondent original z-stack (models 1, 2 and 4) or was assigned randomly from a Gaussian probability distribution (model 3) based on the cFos cell density (**Table 2**) and the total number of cells in the grid, similarly to what above described for the apoptotic cell number.

**Table 2.** Summary of experimental values used for mathematical modelling.

| Experimental value | Group | Mean | SD | n |
| --- | --- | --- | --- | --- |
| Apoptotic/NeuN<sup>+</sup> | **WT** | 0.0005 | 0.0003 | 4 |
|  | **KO** | 0.0052 | 0.0007 | 4 |
| cFos/NeuN<sup>+</sup> | **WT** | 0.0240 | 0.0075 | 4 |
|  | **KO** | 0.0265 | 0.0073 | 4 |

#### 3. Simulations

For each v3D grid and each of the four models, 10,000 randomizations were performed, and the Euclidean distance between each apoptotic cell and its nearest cFos^+^ cell was computed in each randomization. Specifically, we performed 100 cell number (*rn*) randomizations, each followed by 100 cell location *(rl)* randomizations. In each *rm,* the number of cFos^+^ and apoptotic cells in the grid was defined by different criteria according to the four different models detailed above. For each model and for each condition (WT or KO), the cumulative probability distribution of the NN distances between cFos and apoptotic cells was next calculated with 1 μm step and with 99% confidential intervals, by pooling data from the 10,000 simulations and the eight corresponding (WT or KO) v3D grids.

### Statistical analysis

SigmaPlot (Systat Software) was used for statistical analysis. Data were tested for normality and homoscedasticity. Two-sample experiments were analyzed by Student’s t test and more than two-sample experiments by ANOVA. In two-way ANOVA, when interactions between factors were found, the analysis of the relevant variable was split into several one-way ANOVAs and Holm–Sidak method was used as a post hoc. Only p<0.05 is reported to be significant. Data are shown as mean ±SEM.

## RESULTS

### Microglial phagocytosis is uncoupled to apoptosis in *Cstb* KO mice at P30

To determine the apoptosis/phagocytosis dynamics in *Cstb* KO mice we first focused on analyzing hippocampal atrophy and neuronal apoptosis in the hippocampus in young mice at P30, when cerebellar apoptosis and cortical atrophy occur and seizures begin^7,9^ We analyzed the granule cell population by immunostaining the dentate gyrus (DG) of both wild type (WT) and *Cstb* KO mice with the nuclear marker DAPI to assess apoptosis by karyorrhexis and/or pyknosis, and with Iba1 to visualize microglia (**Figure 1A**). We found a significant increase in the total number of apoptotic cells (**Figure 1B**) that was translated into a decrease in the total granule cell population in *Cstb* KO mice compared to WT mice (**Fig 1C**). However, the total granule cell density was similar in both genotypes, with a trend towards a reduction in the volume of the septal hippocampus (p=0.052), (**Fig. 1D, E**). This increased apoptosis and trend to decreased hippocampal volume in *Cstb* KO mice at P30 are consistent with the cerebellar and cortical atrophy previously described^9–11^.

**Figure 1.**
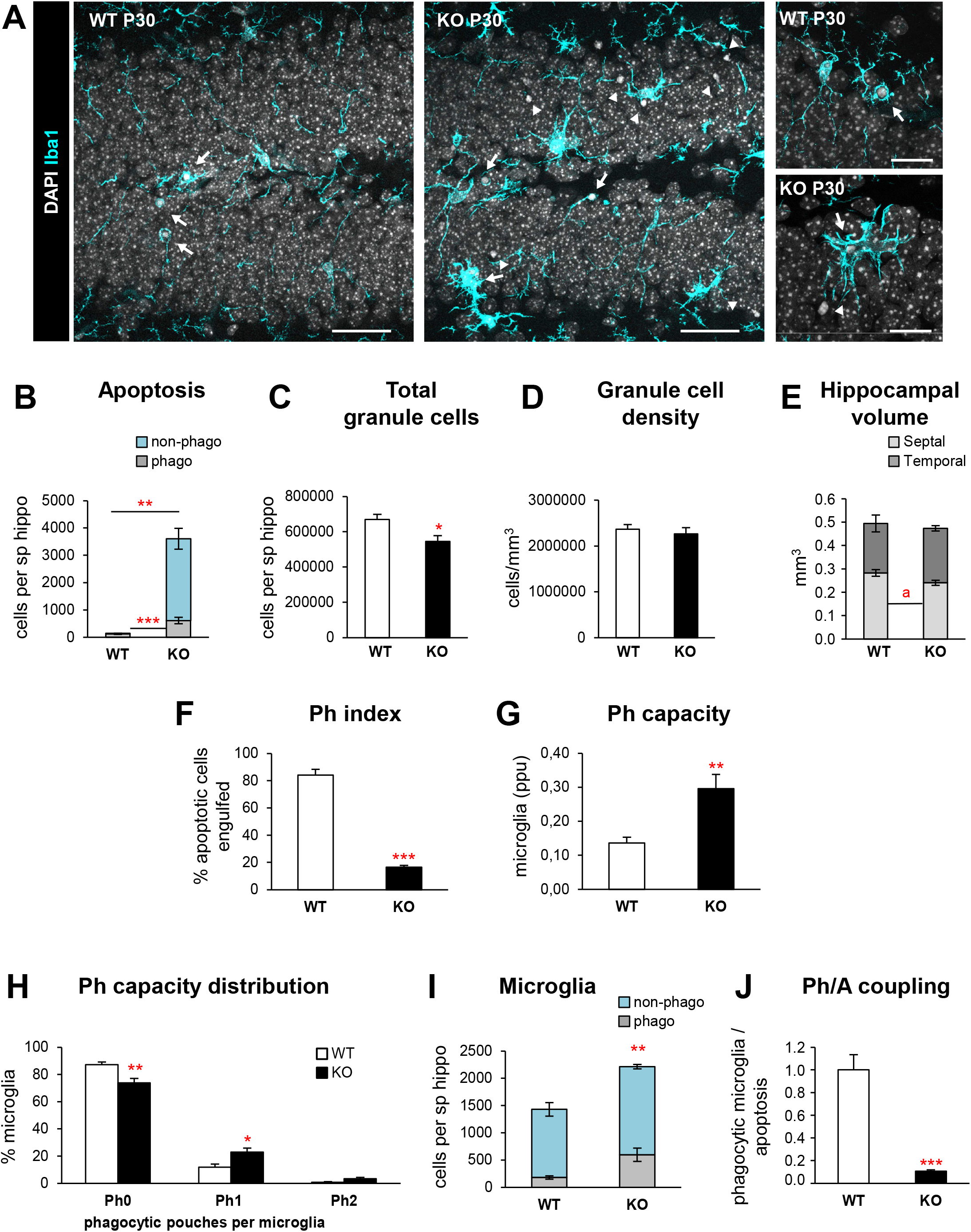
Microglial phagocytosis is impaired in the DG of P30 *Cstb* KO mice. (**A**) Representative confocal images of the DG in WT and *Cstb* KO P30 mice (A). Healthy or apoptotic (pyknotic/karyorrhectic) nuclear morphology was visualized with DAPI (white) and microglia were stained for Iba1 (cyan). High magnification examples of phagocytic microglia following the typical “ball and chain” (upper panel) form with the tip of their processes and phagocytosing with their soma (lower panel). Arrows point to apoptotic cells engulfed by microglia (Iba1+) and arrowheads point to non-phagocytosed apoptotic cells. (**B**) Number of apoptotic cells (pyknotic/karyorrhectic) in the septal hippocampus (n = 6 animals per condition) showing both phagocytosed and non-phagocytosed apoptotic cells. (**C**) Total number of granule cells per septal hippocampus. (**D**) Density of granule cells (per mm^3^) in the septal hippocampus of both WT and *Cstb* KO P30 mice. (**E**) Proportion (in mm^3^) between the volume of the septal and the temporal portions of the hippocampus in WT and *Cstb* KO P30 mice. (**F**) Phagocytic index (in % of apoptotic cells being engulfed by microglia) in the septal hippocampus. (**G**) Weighted phagocytic capacity (Ph capacity) of DG microglia (in parts per unit, ppu). (**H**) Histogram showing the Ph capacity distribution of DG microglia (in % of microglial cells with 0-2 phagocytic pouches, Ph). (**I**) Number of phagocytic (Iba1 + with DAPI inclusions) and non-phagocytic microglial cells (Iba1+ with no DAPI inclusions) per septal hippocampus. (**J**) Ph/A (in fold change) in the septal hippocampus. Bars represent the mean ± SEM. * indicates *p* < 0.05, ** indicates *p* < 0.01, *** indicates *p* < 0.001*, a* indicates *p* = 0.052 by 1-tail Student’s t-test. Scale bars= 40μm (**A**, low magnification), 20 μm (**A**, high magnification); z=7μm (**A**, low magnification), 3.5μm (**A**, high magnification).

Next, we analyzed microglial phagocytosis by measuring the phagocytic index (Ph index), i.e., the percentage of apoptotic cells engulfed by microglia. Microglia in *Cstb* KO mice engulfed 16.6±1.2% of the apoptotic cells, significantly less than WT mice (84.3±4.1%, p<0.001; **Figure 1F**), suggesting that *Cstb* KO microglia was altered. To directly test the proportion of microglial cells that were engaged in phagocytosis, we analyzed their phagocytic capacity (Ph capacity), i.e., the number of phagocytic pouches containing an apoptotic cell per microglia. *Cstb* KO mice showed an increased Ph capacity compared to WT mice, with fewer non-phagocytic cells and more cells with one phagocytic pouch (**Figure 1G, H**). Microglial numbers were also increased in *Cstb* KO mice compared to WT mice (**Figure 1I)**, leading to an increase in net phagocytosis (number of microglia multiplied by the Ph capacity). Nonetheless, this increase in net phagocytosis was insufficient to compensate for the increase in apoptosis, leading to a reduction in the Phagocytosis/Apoptosis coupling (ratio between net phagocytosis and total number of apoptotic cells; **Figure 1J**). Thus, despite the attempt to increase microglial numbers and Ph capacity, microglia were not able to remove the increasing numbers of dead cells in *Cstb* KO at P30.

As we had observed an increase in the total number of microglia (**Figure 1I**), we then analyzed whether this effect was due to microglial proliferation using Ki67, a marker for proliferating cells that is expressed during all active phases of the cell cycle^35^ (**Figure 2 A, B**). We observed a significant increase in proliferating microglia in *Cstb* KO mice compared to WT mice at P30 (**Figure 2C**). In addition, *Cstb* KO microglia presented a hypertrophic morphology (**Figure 1A, 2A**), which was reminiscent of the ameboid-like, multinucleated phenotype of microglia observed in the KA model of epilepsy^15,36^. To assess whether *Cstb* KO microglia also showed multinuclearity we quantified the percentage of cells with more than one nucleus (**Figure 1D**). We found that while multinucleated cells were absent in control mice, they represented 13,1% ±1,5 of microglia in *Cstb* KO mice, similar to what we observed 3d and 7d after KA injection^15^.

**Figure 2.**
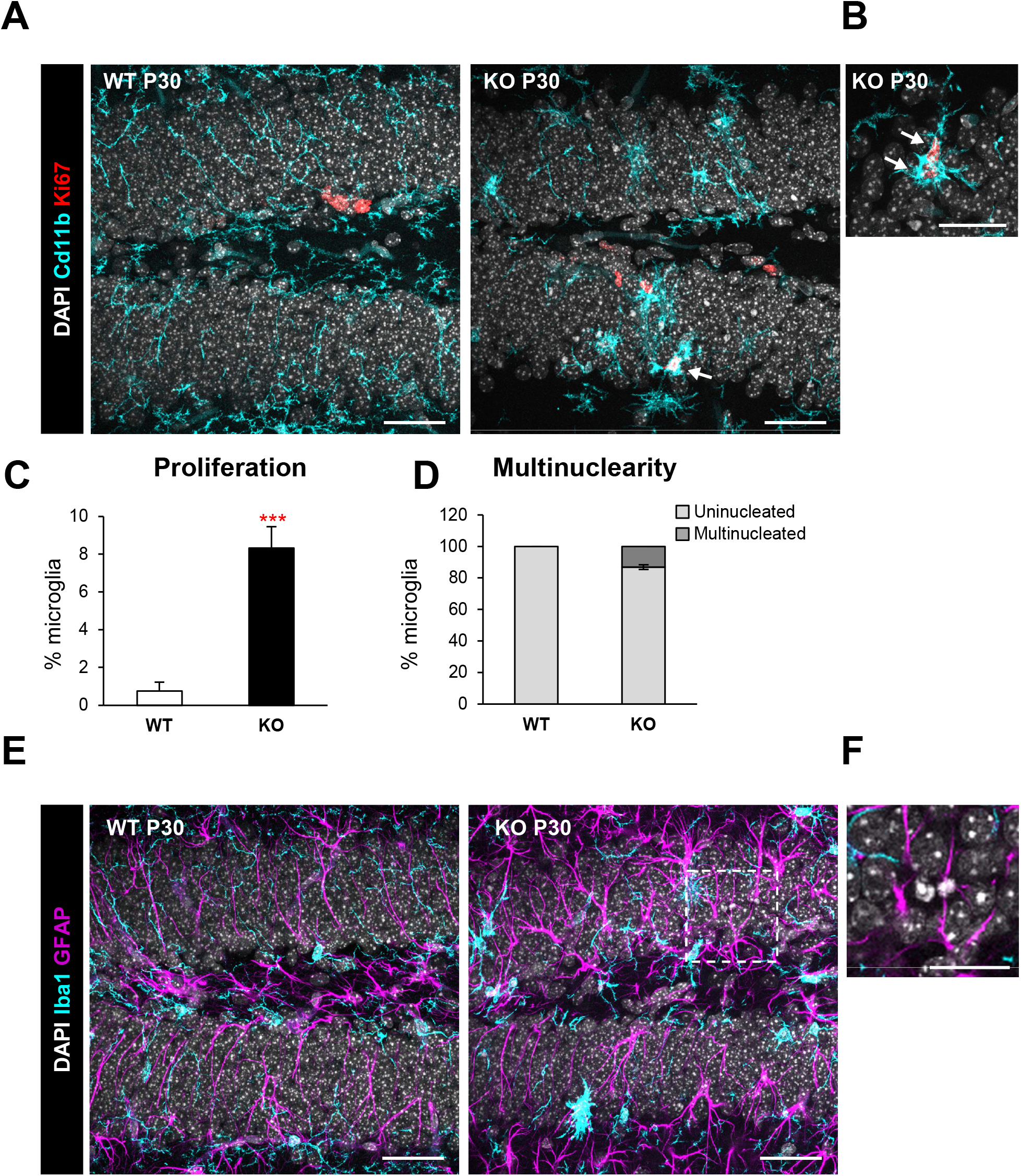
Increased proliferation and multinuclearity in the DG of P30 *Cstb* KO mice. (**A**) DG general view of both WT and *Cstb* KO P30 mice. Proliferating cells were stained for Ki67 (marker for all the active phases of the cell cycle), nuclei (DAPI) and microglia (CD11b). (**B**) High magnification image of a proliferating Ki67+ microglia in P30 *Cstb* KO mice. Arrows point to Ki67+ microglia (**A, B**,). (**C**) Percentage of proliferating microglia assessed by the marker Ki67+. (**D**) Proportion (in %) between uninucleated and multinucleated microglia in the septal hippocampus of WT and *Cstb* KO P30 mice. (**E**) Representative images of the DG in WT and *Cstb* KO mice stained for DAPI (nuclei), microglia (Iba1) and astrocytes (GFAP). (**F**) High magnification image of an apoptotic cell not phagocytosed by neither microglia or astrocytes in *Cstb* KO mice. Bars represent the mean ± SEM. * indicates *p* < 0.05, *** indicates *p* < 0.001 by 1-tail Student’s t-test. Scale bars= 40μm (**A**), 20μm (**B**), 40μm (**E**), 20μm (**F**); z=7 μm (**A**, **E**), 3.5μm (**B**).

Finally, we addressed whether the impairment in microglial phagocytosis was compensated by recruitment of other cell types for phagocytosis. Microglia are specialized phagocytic cells, however, other cell types, such as astrocytes, can also perform phagocytosis under certain conditions^16^. To characterize whether astrocytes contributed to the phagocytosis of apoptotic cells we immunostained the DG for the astrocytic marker GFAP, as well as CD11b for microglia and DAPI for the nuclei (**Figure 2E, F**). We observed a notable morphology change in astrocytes, with cells showing thicker processes, in agreement with previously reported reactive astrogliosis observed in the *Cstb* KO at P30, the onset of myoclonus^11^, but no evidences of their involvement in phagocytosis. Overall, these results demonstrate microglial phagocytosis was impaired and not compensated by astrocytes in *Cstb* KO mice.

### *Cstb* knock-down in microglia does not alter phagocytosis *in vitro*

To determine a possible cell-autonomous effect of *Cstb* on microglial phagocytosis we first addressed whether microglia expressed *Cstb in vivo.* For this purpose we FACS-sorted microglia from P30 fms-EGFP hippocampi (**Figure 3A**), where microglia constitutively express the EGFP protein^25,26^, allowing the discrimination of microglia from other cell types. We analyzed the mRNA expression of *Cstb* as well as downstream *cathepsins B, L* and *S* by RT-qPCR, because *Cstb* mutations are related to increased cysteine protease expression^37^ and activity^38,39^. We found that *Cstb* and *cathepsins B* and *L* were expressed by both microglia and non-microglia cells, whereas *cathepsin S* was solely expressed by microglia (**Figure 3B**). While *Cstb* was not enriched in microglia compared to other cell types, its robust expression suggested that the phagocytosis impairment could be the direct consequence of microglia lacking *Cstb* per se.

**Figure 3.**
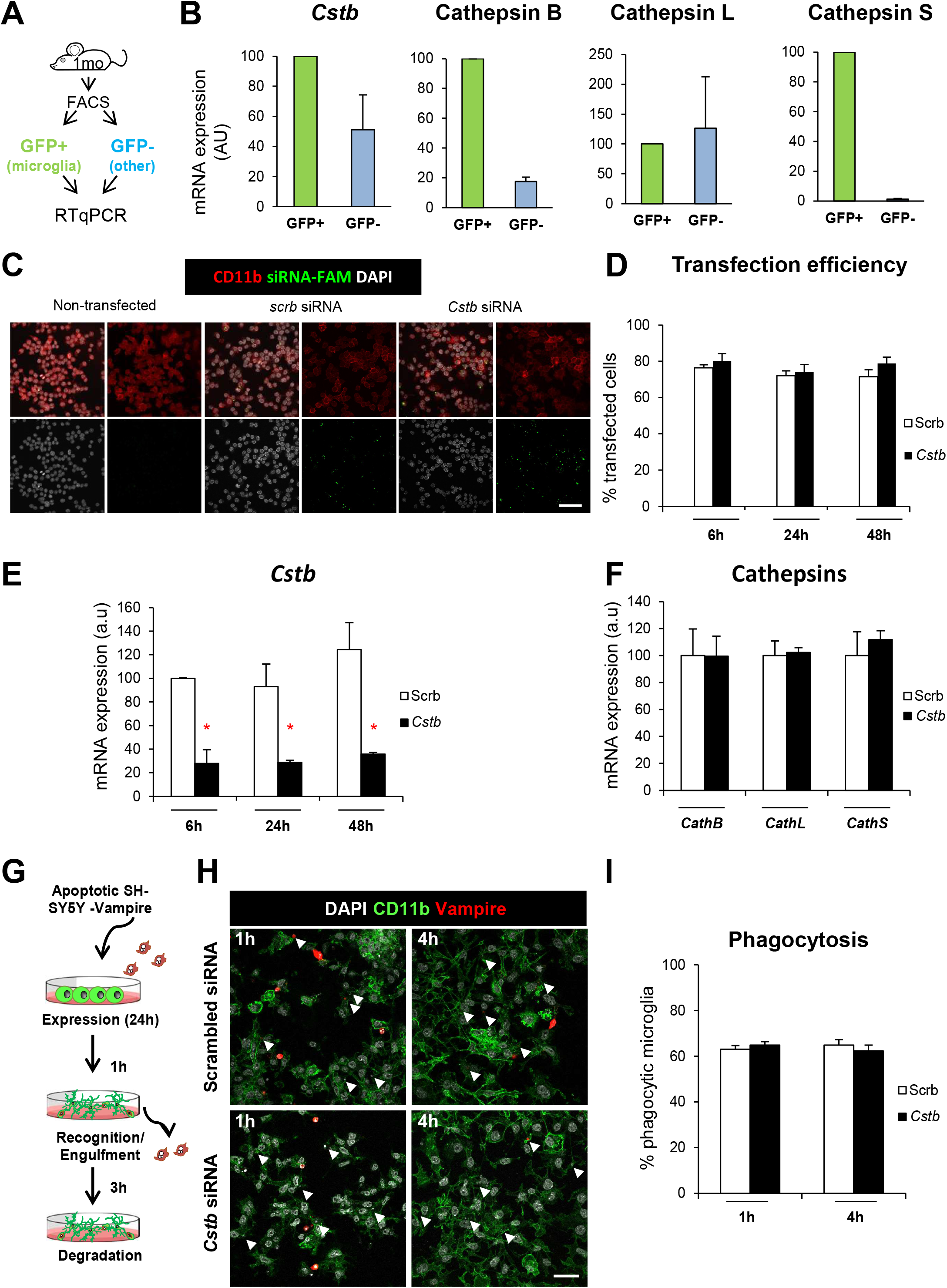
*Cstb* knockdown in microglia does not alter phagocytosis *in vitro.* **A**) Experimental design used to isolate microglia (GFP+) from non-microglial cells (GFP-) for the hippocampi of 1-month-old mice using flow cytometry and RT-qPCR for gene expression analysis. (**B**) Expression of CSTB gene and cathepsins B, L and S in microglia (GFP+) versus non-microglial cells (GFP-) in FACS sorted cells from EGFP-fms mice hippocampi. OAZ1 (ornithine decarboxylase antizyme 1) was selected as a reference gene. (**C**) Representative confocal images of non-transfected (left panels) and scrambled/*Cstb* siRNA transfected BV2 microglia (middle and right panels). Nuclei are stained with DAPI (white), BV2 microglia were stained for CD11b (red) and siRNA transfection was assessed by FAM (green) labeling. (**D**) Percentage of scrambled/*Cstb* siRNA transfected cells along a time course (6, 24 and 48 hours). (**E**) RT-qPCR *Cstb* gene expression in BV2 cells after *Cstb* siRNA silencing through a time course (6, 24 and 48 hours), using OAZ1 as a reference gene. (**F**) RT-qPCR cathepsins B, L, S gene expression in BV2 cells 24 hours after siRNA *Cstb* silencing, using OAZ1 as a reference gene. (**G**) Experimental design of the phagocytosis assay performed 24 hours after BV2 siRNA transfection. Knockdown BV2 cells are fed for 1 and 4 hours with apoptotic SH-SY5Y vampire neurons. (**H**) Representative confocal images of scrambled and *Cstb* siRNA transfected BV2 cells (CD11b staining, green) fed with apoptotic SH-SY5Y vampire neurons (red) for 1 and 4 hours. Arrowheads show phagocytosed SH-SY5Y vampire fragments or full cells. (**I**) Percentage of phagocytic BV2 cells after 1 and 4 hours of phagocytosis. Only particles fully enclosed by BV2 pouches were identified as phagocytosis. Bars represent the mean ± SEM. * indicates *p* < 0.05, *** indicates *p* < 0.001 by two-way ANOVA. Scale bars= 60 μm (**C**), 40 μm (**H**).

To directly assess the effect of microglial *Cstb* on phagocytosis, we set up an *in vitro* model of *Cstb* knock-down in the microglial cell line BV2 (**Fig 3C-I**). We transfected BV2 microglia with 6-carboxyfluorescein (FAM) labeled siRNAs against *Cstb* or a scrambled siRNA as a control (**Figure 3C**). We obtained a high transfection efficiency through a time course of 6, 24 and 48h (**Figure 3D**) and validated the transcription down-regulation of the *Cstb* gene by RT-qPCR through the time course. We observed that the expression of *Cstb* was greatly reduced up to 48 hours (**Figure 3E**), whereas the expression of the related cathepsins was not affected by *Cstb* siRNA treatment (**Figure 3F**). Control and *Cstb* KO microglia were then fed with apoptotic SHSY5Y-vampire neurons for 1 and 4h, in which apoptosis was previously induced (Staurosporine 3μM, 4 hours) ^15^ (**Figure 3G**). Microglia were stained with CD11b, apoptotic neurons constitutively expressed the red fluorescent protein and nuclei were stained with DAPI (**Figure 3H**). We found no differences in phagocytosis in microglia treated with either scrambled or *Cstb* siRNA (**Figure 3I**). Our *in vitro* model, nonetheless, does not fully recapitulate the *in vivo* situation, because *Cstb* KO mice have *Cstb* chronic depletion that is accompanied by modulation of cathepsin activity and/or expression^37,39^. Nonetheless, these results suggest that the cell-autonomous acute *Cstb* deficiency in microglia is not sufficient to induce the phagocytosis impairment observed *in vivo.* We, therefore, searched for alternative mechanisms to explain the reduced phagocytosis in *Cstb* KO mice. As we had previously shown that seizures interfere with phagocytosis in a mouse model of MTLE, our next step was to determine if the phagocytosis blockage was related to seizures by analyzing an early developmental stage, P14.

### Phagocytosis impairment is specific of the granule cell layer in cstb KO mice at P14

At P14, *Cstb* KO mice are asymptomatic and do not have seizures^11^ and in agreement we did not observe any changes in the number of active neurons, labelled with the immediate early gene cFos, whose expression is rapidly induced upon depolarization^40^ (**Figure 4A-C**). cFos^+^ neurons were found at similar numbers in both WT and *Cstb* KO mice at P14, but in *Cstb* KO mice they were located exclusively in the GL and not in the SGZ, where radial neural stem cells and their immature progeny reside (**Figure 4A, C**). However, we did notice that whereas in WT mice most apoptotic cells were found in the SGZ, in *Cstb* KO mice apoptotic cells were found mostly in the GL and in very close proximity to cFos^+^ neurons (**Figure 4B**), prompting us to analyze separately GL and SGZ apoptosis and phagocytosis.

**Figure 4.**
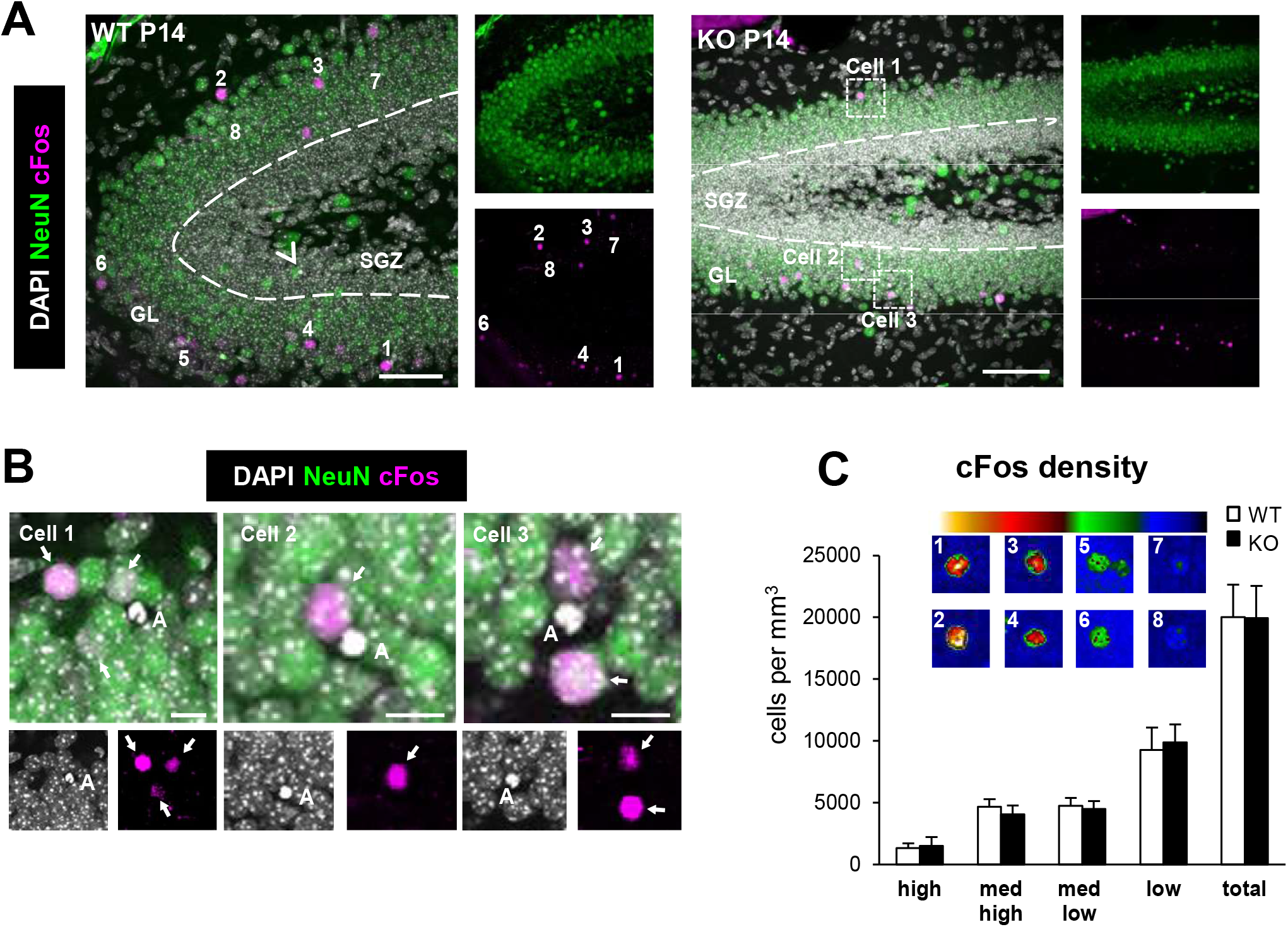
cFos+ cells in the DG of WT and *Cstb* KO P14 mice. (**A**) Representative confocal images of the DG of WT and *Cstb* KO mice. Healthy or apoptotic (pyknotic/karyorrhectic) nuclear morphology was visualized with DAPI (white), neurons were identified with the neuronal marker NeuN (green) and activated neurons were stained for the early expression gene cFos (magenta). Arrowhead points to an apoptotic cell in the SGZ in WT mice; framed apoptotic cells in *Cstb* KO mice are shown in **B**. Numbered cFos^+^ cells are shown in **C**, as cells with high (1,2), medium high (3, 4), medium low (5, 6) and low (7,8) cFos intensity. (**B**) High magnification examples showing the close proximity between apoptotic cells (DAPI, white) and cFos+ neurons (magenta) in Cstb KO mice. Granular neurons are stained with NeuN (green). Arrows point to cFos+ neurons. (**C**) Distribution of cFos+ cells in WT and *Cstb* KO mice (per mm^3^). The color code indicates the classification criteria of the cFos+ cells based on their intensity (high, medium high, medium low, low). A total of 1046 cells for WT P14 mice and 713 cells for *Cstb* KO mice were quantified and classified according to their cFos expression. No significant differences were found. Scale bars= 50μm (**A**), 10μm (**B**); z=28 μm (**A**, WT), 17.5 μm (**A**, KO).

In the SGZ, we did not observe any significant difference in apoptosis nor phagocytosis between WT and *Cstb* KO mice at P14 (**Figure 5A-F**). In contrast, in the GL there were few apoptotic cells in WT mice and a 10-fold increase in *Cstb* KO mice (**Figure 5D, E**).

**Figure 5.**
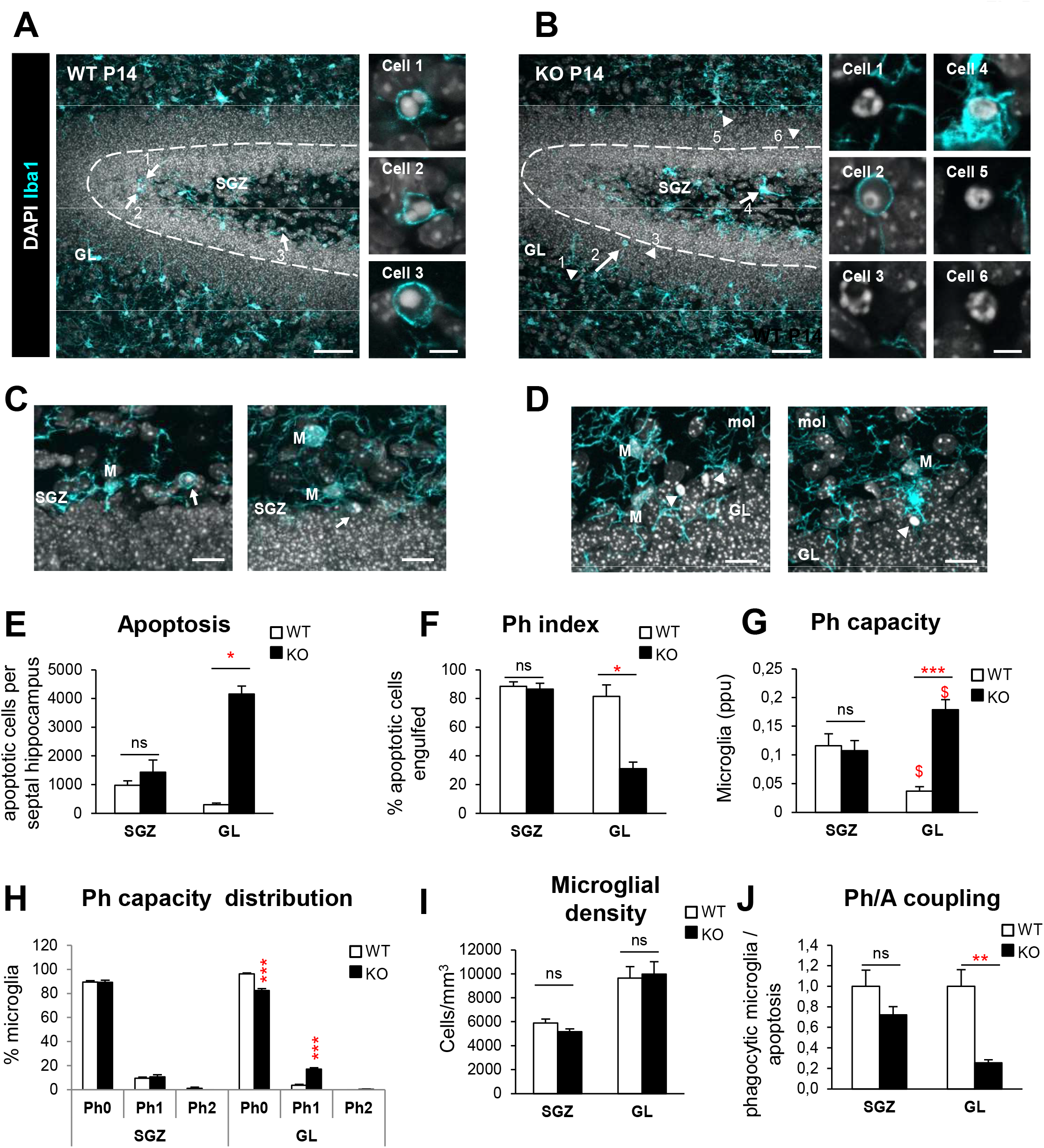
Phagocytosis impairment is specific of the granule cell layer in *Cstb* KO mice at P14. (**A, B**) DG general view of both WT and *Cstb* KO P14 mice, nuclei are stained with DAPI (white) and microglia with Iba1 (cyan). Close up images show phagocytosed and non-phagocytosed apoptotic cells in WT and *Cstb* KO P14 mice. (**C, D**) Representative images of apoptotic cells (condensed DAPI) engulfed by microglia (M, cyan) in the SGZ of WT P14 mice (**C**) and non-phagocytosed cells in the GL of *Cstb* KO P14 mice (**D**). Arrows point at phagocytosed apoptotic cells and arrow heads to non-phagocytosed apoptotic cells (**A-D**). (**E**) Number of apoptotic cells (pyknotic/karyorrhectic) both in the SGZ and GL, per septal hippocampus (n =12 animal for each condition). (**F**) Phagocytic index (in % of apoptotic cells being engulfed by microglia) in the SGZ and GL of the septal hippocampus in WT and *Cstb* KO P14 mice. (**G**) Histogram showing the Ph capacity distribution of DG microglia (in % of microglial cells) in the SGZ and GL. (**H**) Microglial density (cells/mm^3^) per septal hippocampus both in WT and *Cstb* KO P14 mice, distinguishing between SGZ and GL. (**I**) Weighted Ph capacity of DG microglia (in ppu). (**J**) Ph/A (in fold change) in the SGZ and GL of the septal hippocampus in WT and *Cstb* KO P14 mice. Bars represent the mean ± SEM. * indicates *p* < 0.05, ** indicates *p* < 0.01, *** indicates *p* < 0.001 by Student’s t-Test comparing WT vs KO. Scale bars= 50μm (**A, B**), 5μm (inserts in **A, B**), 30μm (**C, D**); z=18.9 μm (**A,B**), 9.8 μm (**C left**), 16.1 (**C right**), 11.2 μm (**D left**), 12.6 μm (**D right**).

The majority of these GL apoptotic cells were not phagocytosed, resulting in a reduced GL Ph index in cstb KO mice compared to WT mice, at P14 (**Figure 5F**). Similar to the global effect we had observed before in the whole dentate gyrus at P30, we found that GL microglia increased their Ph capacity in *Cstb* KO mice compared to WT mice although this effect was insufficient to cope with the increased number of apoptotic cells (**Figure 5G, H**). In this early stage, we found no obvious morphological changes nor increases in microglial numbers (**Figure 5I**). Overall, we found that the increase in apoptotic cells was not compensated by a sufficient increase in phagocytosis, resulting in an uncoupling between apoptosis and phagocytosis (**Figure 5J)**. Overall, these results demonstrate that the microglial phagocytosis impairment was specific of the GL and preceded seizure development in *Cstb* KO mice at P14. This data suggested that there could be functional differences between GL and SGZ that could explain the GL-specific increase in apoptosis and microglial phagocytosis impairment.

### Apoptotic cells are in close proximity to active cFos^+^ neurons

The close vicinity of apoptotic cells to GL cFos^+^ neurons in P14 mice (**Figure 4B**), suggested that the phagocytosis efficiency of GL microglia could be related to neuronal activity, as neuronal hyperactivity during seizures prevents microglia from targeting apoptotic cells^15^. To address whether the proximity of cFos^+^ neurons had an impact on apoptosis and phagocytosis, we directly estimated the distance to each phagocytosed and non-phagocytosed cell to the closest cFos^+^ neuron (nearest neighbor, NN) (**Figure 6A**). Because of the limitation imposed by the relatively small thickness of our z-stacks (border effect; see Methods for more detail), we only had a sufficient number of apoptotic cells in KO mice (n=15 phagocytosed and n=5 non-phagocytosed apoptotic cells from n=9 mice, from an initial set of 295 cells).

**Figure 6.**
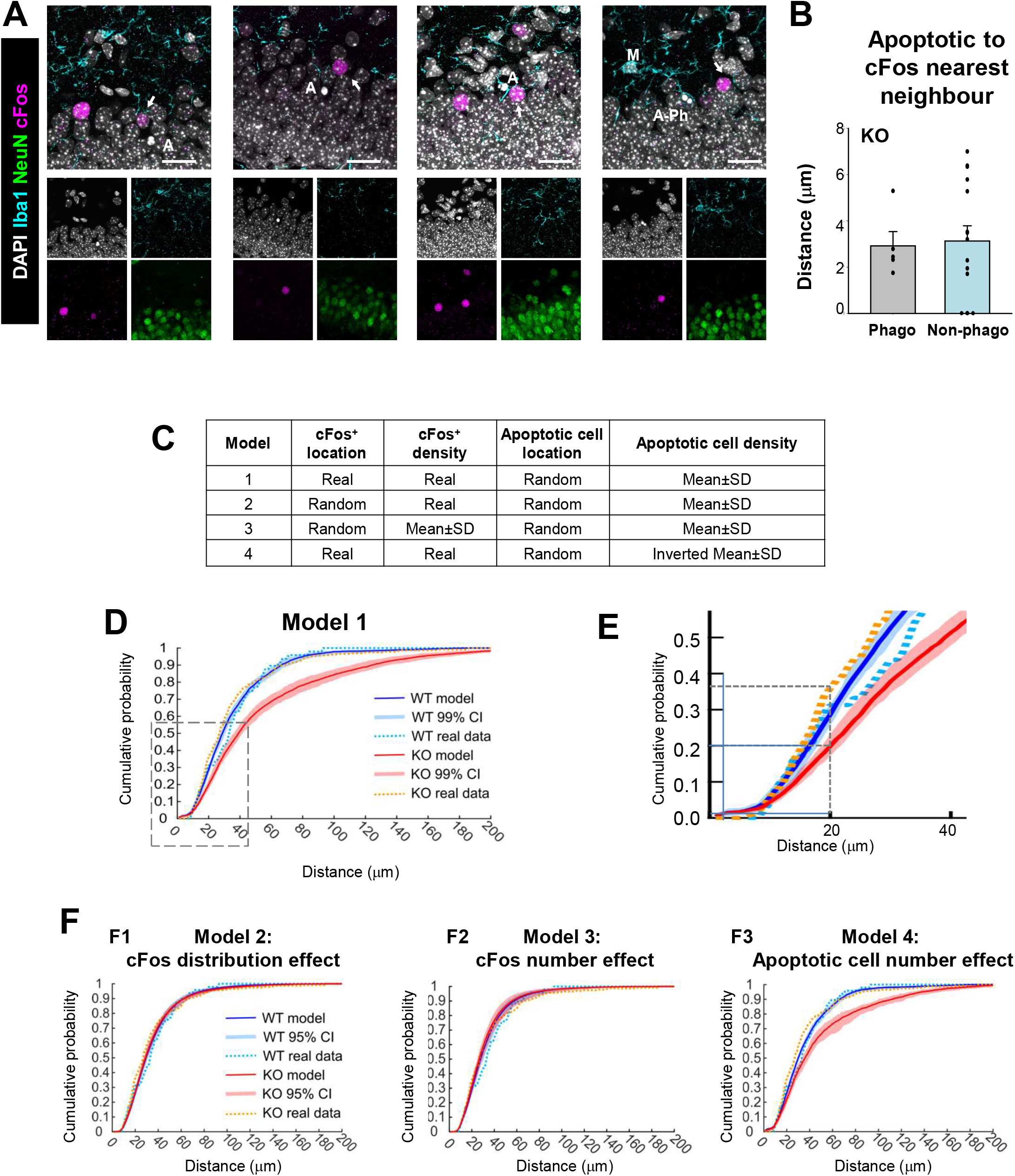
Proximal relationship between apoptotic cells and cFos+ neurons in the granule cell layer of *Cstb* KO mice. (**A**) Representative confocal images of the GL of *Cstb* KO mice. Healthy or apoptotic (pyknotic/karyorrhectic) nuclear morphology was visualized with DAPI (white), neurons were identified with the neuronal marker NeuN (green), microglia with Iba1 (cyan) and activated neurons were stained for the early expression gene cFos (magenta). Both phagocytosed (A-Ph) and non-phagocytosed apoptotic cells (A) were close to cFos^+^ neurons (arrows). (**B**) Quantification of distance from phagocytosed and non-phagocytosed apoptotic cells to the cFos+ NN, for those cells that met the inclusion criteria (see Methods). (**C**) Summary of the different simulation models based on the location and density of cFos^+^ and apoptotic cells. (**D**) Cumulative probability of the distances between apoptotic cells and NN cFos^+^ neurons for WT (blue) and *Cstb* KO mice (red), resulting from 10,000 simulations of a virtual 3D model, indicating the 99% confidence interval (CI). The cumulative probability for real (measured) data is shown for WT (light blue) and *Cstb* KO (orange). (**E**) Amplification of the area showed in C. (**F**) Cumulative probabilities of the distances between apoptotic cells and NN cFos^+^ neurons for WT (blue) and *Cstb* KO mice (red), resulting from 10,000 simulations of the indicated virtual 3D model. Scale bars indicate 20μm. z= 9.1 μm, 7.7 μm, 16.8 μm, 9.1 μm (**A**, from left to right).

In *Cstb* KO mice, both phagocytosed and non-phagocytosed cells were found very close to the cFos^+^ NN (3.0 ± 0.5 and 3.0 ± 0.6μm, respectively; non-significant difference; **Figure 6B**). To determine whether this short distance would be expected if cells were homogenously distributed through the thickness of our z-stacks, we calculate that 50% of the cells should be located at ¼ distance to either z-border (green-shaded area in **Supplementary Figure 1**), that is, at 5.9 ± 0.2 and 6.0 ± 0.1μm to the cFos^+^ NN for phagocytosed and non-phagocytosed cells, respectively (**Supplementary Figure 1)**. Thus, both phagocytosed and non-phagocytosed apoptotic cells in *Cstb* KO mice seem to be much closer (3μm) to active neurons than would be expected. Although we could not detect significant differences between phagocytosed and non-phagocytosed cells in *Cstb* KO mice, these results do suggest an unexpected relationship between neuronal activity and apoptosis.

To strengthen the analysis of distance between apoptotic cells and NN cFos^+^ neurons and in both WT and KO mice and overcome the limitation of analyzing only 20 cells, we developed a mathematical model (see Methods for details). We created a virtual 3D model of the GL based on the spatial distribution and density of neurons, cFos^+^ neurons and apoptotic cells (**Supplementary Figure 1**). In this model we reproduce the same experimental limitations imposed by the limited thickness of the z-stacks (see Methods) compared the size of the (XY) field of view and therefore no cells were discarded due to the border effect. Using this virtual 3D model we compared the entire experimental distribution of NN distance between cFos^+^ and apoptotic cells to the distribution generated by the 10,000 simulations. Specifically, we developed four different models (**Figure 6C**).

In the first model cFos^+^ neurons were located in their original position (in the z-stack) and the position of the apoptotic cells was randomized, and their numbers were extrapolated from the experimental cell density (**Figure 6C**). For each apoptotic cell we performed 10,000 randomizations and calculated the distance to the NN cFos^+^ cell. Finally, the cumulative probability to encounter a cFos^+^ cell within a given distance from an apoptotic cell was computed with its 99% confidence interval both in the WT- and KO-derived v3D grids (**Figure 6D, E**). In WT mice, the measured apoptotic-cFos^+^ NN distance was within the modeled confidence interval up to a distance of 22μm, which matches the average z-stack thickness (24.2 ±0.4μm) and validates that our model is isotropic, i.e., does not depend on the orientation and therefore is within the range of the z-stack thickness. In contrast, in *Cstb* KO mice the measured percentage of NN cFos^+^-apoptotic cells between 17 and 24 μm was significantly higher than what observed both in the WT and *Cstb* KO models. For instance, in the *Cstb* KO model 20% of the apoptotic cells are found at less than 20μm distance from the cFos^+^ NN, whereas in the real data from *Cstb* KO mice, 20% of the apoptotic cells were found at less than 16μm and 35% of the apoptotic cells are found at less than 20μm (**Figure 6E**). This data suggested that as in our direct quantifications (**Figure 6B**), apoptotic cells were closer than expected to active cFos^+^ neurons in *Cstb* KO mice.

Nonetheless, the modeled distance was significantly larger in KO than in WT mice (**Figure 6E**). To further understand the role of other parameters, specifically cell number or location, on the model, we used other three different models (**Figure 6C**): Model 2 to test the impact of cFos^+^ cell location, Model 3 to test cFos^+^ cell density, and Model 4 to test apoptotic cell density. We first tested the effect of cFos^+^ cell location (Model 2), by randomly positioning both cFos^+^ neurons and apoptotic cells (**Figure 6F1**). In this model, there were no differences between the modeled NN distances between WT and KO. Importantly, in KO mice the difference between the measured and modeled apoptotic-cFos^+^ NN distance was strongly reduced although it was still significant with a maximum around 20μm. The reduced difference between the measured and modeled NN distance in KO mice in model 2 suggest that the differences between the modeled NN distances in Model 1 originated from a differential distribution of cFos^+^ cells between WT and KO.

We then tested the effect of cFos^+^ density by creating a Model 3 in which cFos^+^ cells were positioned randomly in the v3D grid and their number was estimated from the experimentally measured cFos density rescaled by the number of cells in the grid, whereas apoptotic cells were modeled randomly as above (**Figure 6F2**). The results of Model 3 were similar to Model 2, with no differences between modeled and measured NN distances for WT and KO, suggesting that the number of cFos^+^ cells was not related to the differential effect found in Model 1. Finally, we tested the effect of different density of apoptotic cells in the WT and KO modeled distances in Model 4. For this, we went back to use the real distribution of cFos+ cells (as in Model 1) and positioned the apoptotic cells randomly but with an inverted density: WT model using KO apoptotic cell density and viceversa (**Figure 6F3**). The results of Model 4 were similar to Model 1: the modeled NN distance was larger in KO than in WT, and apoptotic cells were found closer to active neurons than expected in KO mice. Overall, this data suggests that while *Cstb* KO mice have similar cFos^+^ density and intensity at P14 (**Figure 4C**) and do not have yet seizures^11^, they have an abnormal distribution of cFos^+^ active neurons that results in a closer distance to apoptotic cells (**Figure 6B,C**). These results suggest a very close relationship between abnormal neuronal activity in *Cstb* KO mice and apoptotic cells. Indirectly, they also suggest a relationship with the phagocytosis impairment found in Ctsb KO mice, because the number of apoptotic cells observed is the net result between apoptosis induction and phagocytosis removal.

## DISCUSSION

In this paper, we show for the first time that microglial phagocytosis of apoptotic cells is impaired in *Cstb* KO mice, a model that recapitulates the main features of clinical EPM1. Additionally, we provide evidences on the possible mechanisms that underlie the phagocytic disruption associated with CSTB deficiency, based on the following findings: First, microglial phagocytosis of apoptotic cells was reduced in the hippocampus of symptomatic *Cstb* KO mice, at P30. Second, the cell-autonomous lack of *Cstb* in microglia did not drive phagocytosis impairment, since *Cstb* down-regulation in pure microglial cultures did not alter phagocytosis efficiency. Third, microglial phagocytosis impairment was already present in the hippocampus of *Cstb* KO mice at P14, prior to the onset of seizures, suggesting that microglial phagocytosis impairment appears at early stages of disease and independent of seizures. Fourth, the impairment was specific for the GL, whereas the neurogenic niche of the SGZ was spared. Fifth, a virtual 3D model of the hippocampal GL in young CSTB KO mice (P14) predicted an aberrant distribution of cFos active neurons that results in a closer distance to apoptotic cells. These results suggest that local neuronal activity may alter apoptosis dynamics in *Cstb* KO mice, which may explain, at least in part, the reported impairment on microglial phagocytosis associated with CSTB deficiency. Here, we will first discuss the pathological effects of *Cstb* deficiency in the hippocampus, including atrophy, apoptosis and phagocytosis impairment. Then we will examine possible mechanisms underlying this impairment, including a cell-autonomous effect of *Cstb* on microglia and environment-related factors such as seizures and local neuronal activity.

### Early symptomatic *Cstb* KO mice exhibit slight atrophy of the hippocampus

EPM1 is the most common type of progressive myoclonus epilepsy, a heterogeneous group of inherited diseases that concur with myoclonus, epilepsy, and progressive neuronal degeneration^41^. Most EPM1 patients are homozygous for a promoter region repeat expansion mutation resulting in significantly reduced CSTB expression and approximately 10% of CSTB expression left in their cells^42^. Patients with total lack of CSTB display an early infantile onset rapidly progressing encephalopathy^43,44^. Progressive loss of brain volume that affects different structures has been reported in patients with CSTB mutations^5,6,44,45^ as well as in *Cstb* KO mice^3,10,11,46^. At early disease stages in *Cstb* KO mice, apoptosis of granule neurons^7^ and atrophy^11^ are most prevalent in the cerebellum. Here we show similar effects in the hippocampus, as young early symptomatic (P30) *Cstb* KO mice contained fewer hippocampal granule cells and a tendency to atrophy of the septal hippocampus, with no changes at the hippocampal temporal region. In agreement, an age-dependent hippocampal volume loss was reported in *Cstb* KO mice using *in vivo* magnetic resonance imaging (MRI) volumetry starting at 2 months of age and peaking at 6^46^. The differences observed in the initiation of hippocampal atrophy may be explained by the higher sensitivity of stereology based cell quantification methods when compared to MRI structural change assessment^47^. Thus, granule cell death and hippocampal atrophy start early in the brains of *Cstb* KO mice and exhibit a progressive nature.

### Microglial phagocytosis of apoptotic cells is impaired in early symptomatic *Cstb* KO mice

In addition to the neuronal damage, other cell types have also been involved in the pathology of EPM1. Some of the earliest changes pointed towards an altered inflammatory response by microglia, the brain resident macrophages^11,13^. Here we focused on another aspect of microglia: their phagocytic function. Microglia are very efficient phagocytes in the adult hippocampus in physiological conditions, where unchallenged microglia rapidly clears the excess newborn cells produced in the neurogenic niche^22^, actively participating in the regulation of neurogenesis^17^. After stressful stimuli such as inflammation and excitotoxicity, microglia use different strategies to enhance phagocytosis and match the increased apoptosis levels^15^: recruiting more cells to become phagocytic (theoretically, up to 100%), increasing the phagocytic capacity of each microglia (at least up to seven pouches per cell), and proliferating. Combined, these strategies make microglia a very powerful phagocyte.

However, the phagocytic potential of microglia was not fully summoned in the hippocampus of young adult *Cstb* KO mice, at the age when they start to manifest clinical myoclonus (P30). Although microglia tried to compensate for increased apoptosis by 1) increasing the number of apoptotic cells cleared by each microglial cell, and 2) increasing their numbers through proliferation, net phagocytosis did not match increased apoptosis in *Cstb* KO mice. These results are in line with previous mouse and human data of MTLE, where microglial phagocytosis was blocked, leading to accumulation of apoptotic cells in the hippocampus and chronic inflammation^15^. In addition to the phagocytosis impairment, microglia exhibited a hypertrophic morphology and showed multinuclearity in *Cstb* KO mice, suggestive of microglial dysfunction and inflammation^15,48^. These results are also in agreement with the early abnormal morphology and expression of the inflammatory protein F4/80 in microglia prior to gross neurodegeneration in *Cstb* KO mice^11^. Overall, we here demonstrate that the constitutive lack of functional CSTB in mice induces an early pathological microglial status and a reduced clearance of apoptotic cells.

### Microglial phagocytosis disruption is not due to the cell autonomous lack of CSTB

The microglial phagocytosis impairment could be mechanistically related to the lack of *Cstb* in microglia and/or in other brain cell types. Indeed, we here show that *Cstb* expression was not restricted to microglia in the brain, suggesting that cell autonomous (lack of *Cstb* in microglia) or environment-driven (lack of *Cstb* in other cell types) mechanisms could underlie the phagocytosis impairment. However, we observed no alterations in an *in vitro* model of phagocytosis using siRNA to deplete microglial *Cstb.* Nonetheless, this *in vitro* model does not fully recapitulate the full extent of CSTB deficiency in EPM1 patients, which includes a complex modulation of activity and mRNA expression of target cathepsins of CSTB^38,39^. Neither does it fully mimic the complexity of in vivo recognition and engulfment of apoptotic cells, which requires the release of “find-me” signals from apoptotic cells and microglial process motility to engulf the cells (whereas *in vitro* apoptotic cells are simply dumped on top of microglia)^27^. Despite its limitations this data suggests that the lack of *Cstb* in microglia *per se* does not impact the clearance of apoptotic cells and suggests that the phagocytosis impairment could be related to environmental factors associated with the lack of *Cstb,* including its main pathological feature, seizures.

### Microglial phagocytosis impairment is independent of seizure-activity

Neuronal hyperactivity during seizures alters the microglial phagocytic response to apoptotic cells. For example, seizure-induced widespread release of ATP in MTLE masks the microgradients of the “find-me” signal ATP used by microglia to find apoptotic cells^15^. To analyze the impact of seizures in microglial phagocytosis, we took advantage of young *Cstb* KO mice (P14), which do not present clinical seizures. Using cFos as a marker of neuronal depolarization^40^ we did not find significant changes in the number of cFos^+^ neurons but, nonetheless, our virtual 3D model indirectly suggested that they were abnormally distributed in *Cstb* KO mice at P14, which is likely reflected in the seizures that appear at later stages. Interestingly, cFos active neurons, increased numbers of apoptotic cells, and microglial phagocytosis impairment were selectively observed in the GL of the hippocampus and spared the SGZ, illuminating an unexpected relationship between neuronal activity cell death and microglial phagocytosis.

### Local neuronal activity in the GL may contribute to the microglial phagocytosis impairment

In the GL, apoptotic cells were frequently positioned near active neurons labeled with cFos in *Cstb* KO mice, at an average distance of 3μm. In agreement, our virtual 3D model suggested that apoptotic cells were located closer to the active neurons than would be expected based on their relative cell densities. The number of apoptotic cells in a given time point is the net result between apoptosis induction (input) minus phagocytosis (output), and thus these results indirectly suggest suggest that local neuronal activity in the GL could alter apoptosis and phagocytosis dynamics in *Cstb* KO mice. Indeed, neuronal activity dependent molecules such as ATP, exert their functions locally due to limited diffusion to short distances in the tortuous brain parenchyma^50^. For instance, the effective diffusivity of a small molecule such as sucrose (0.342KDa, close to the 0.551KDa of ATP) is 310μm/s, whereas that of larger molecules such as NGF (nerve growth factor, 26.5KDa) is 2.95μm/s. Therefore, the average 3μm distance between apoptotic cells and active cFos neurons found in *Cstb* KO mice implies that the dead cells are well within the influence area of the active neurons’ soma. Thus, both the dead cells and the reaching microglial processes could be affected by the somatic release of modulators by granule neurons, or by the perisynaptic release of modulators by incoming fibers from several afferent systems^51^.

The close relationship between apoptotic cells and active neurons in *Cstb* KO mice is supported by both direct measurements and the virtual 3D model but does not provide evidence that this relationship affects phagocytosis. Nonetheless, it is important to note that the net apoptosis observed is in part the result of the phagocytosis dynamics: the number of apoptotic cells at any given time depends as much on the input (apoptosis induction) as on the output (removal by phagocytosis)^14^. Therefore, our data shows an overall impairment of microglial phagocytosis in the GL of *Cstb* KO mice and suggests a complex scenario in which sub-seizure local neuronal activity may affect the clearance of apoptotic cells by microglia.

In addition, other pathophysiological mechanisms may participate in the phagocytosis impairment observed in *Cstb* KO mice. For instance, one possibility is that intrinsic differences between GL and SGZ microglia explain why the impairment is restricted to the GL. SGZ microglia are likely “trained” for phagocytosis, as they have been exposed to apoptotic cells during the postnatal period, whereas GL microglia is not. This “training” could result in differences in their phagocytosis potential. For instance, SGZ microglia can reach a Ph capacity around 0.8ppu in an LPS model at P30^22^. This number is very far from the 0.18ppu of GL microglia in *Cstb* KO mice at P14 reported here. Indeed, microglial supbopulations in the CA region of the hippocampus have been recently observed upon seizures induced by pilocarpine, based on the expression of keratan sulfate polysaccharides (epitope 5D4), whose overexpression is related to a higher ex vivo phagocytosis of zymosan particles possibly related to the engulfment of synapses^52^. Another potential candidate is the complement system, which is used by microglia to facilitate recognition (opsonization) of apoptotic cells^28^ as well as synapses^53^, and is progressively activated in rodent and human epilepsy^54,55^, including *Cstb* KO mice ^37,56^. Nonetheless, it is beyond the scope of this paper to test the role of complement or keratan sulfate and future studies will decipher the underlying mechanisms of microglial phagocytosis impairment in *Cstb* KO mice.

In summary, we here extend our initial observations that microglial phagocytosis impairment is an early feature of mouse and human MTLE and show that it also occurs in a genetic model of EMP1 by *Cstb* deficiency. We also provide an unexpected link between phagocytosis impairment, accumulation of apoptotic cells and local neuronal activity in these mice that further supports the suggestion that both abnormal local neuronal activity (in presymptomatic *Cstb* KO mice) and network hyperactivity (in MTLE mice) regulate the efficiency of microglial phagocytosis. Phagocytosis is an essential component of the brain regenerative response and therefore, future therapies for epilepsy patients should be aimed not only at reducing neuronal death but also at harnessing microglial phagocytosis.

---

**Supplementary Figure 1. Physical constraints in estimating cFos^+^ to apoptotic cell NN distances.** (**A**) NN distances for WT (white squares) and *Cstb* KO (black squares) in the 25 and 253 apoptotic cells collected from 2 series of sections from the GL of wild-type (WT) and KO mice, compared to the distance of each cell to the closest z-border of the corresponding z-stack. (**B**) Higher magnification of the first 15mm shown in (**A**). The grey line marks the inclusion criteria for the study, where only the cells whose NN distance was smaller to the distance to the nearest z-border were included. Within the inclusion area (dotted line), 50% of the cells would be expected to be found within a distance of 50% the thickness of the z-stack (green-shaded area), i.e., at 25% distance to either the upper or the lower z-stack border, although in fact all cells were found within this region. (**C**) In *Cstb* KO mice the 50% distance was similar between phagocytosed and non-phagocytosed apoptotic cells. (**D**), Virtual model for NN distances. First, a grid was created based on the position of NeuN^+^ neurons in the GL (left panel). On this grid, we positioned cFos^+^ neurons and apoptotic cells in four different models (**Figure 6C**) and calculated the distances of each apoptotic cells to all the cFos^+^ neurons on the grid, to determine the NN. Scale bar= 20 μm (**D**).

## ACKNOWLEDGEMENTS

This work was supported by grants from the Spanish Ministry of Economy and Competitiveness (http://www.mineco.gob.es) with FEDER funds to A.S. (RTI2018-099267-B-I00, BFU2012-32089 and RYC-2013-12817) to A.S. and J.V. (BFU2015-66689) and to P.B (SAF2015-69484-R); a Leonardo Award from the BBVA Foundation to A.S. (IN16,_BBM_BAS_0260); a Tatiana Foundation project to A.S. (P-048-FTPGB 2018); a Basque Government Department of Education project to A.S. (PI_2016_1_0011; http://www.euskadi.eus/basque-government/department-education/); Ikerbasque start-up funds to J.V. and P.B.; and by Folkhälsan Research Foundation (to A.-E.L.) In addition, V.S.-T. and OA are recipients of pre-doctoral fellowship from the Basque Government; S.B. is recipient of pre-doctoral fellowship from the Spanish Ministry of University of the Basque Country EHU/UPV. A.-E.L. is a HiLIFE Fellow at the University of Helsinki. The funders had no role in study design, data collection and analysis, decision to publish, or preparation of the manuscript. We thank Victor Sánchez Zafra for technical support. We are very grateful to Eva Benito for thoroughly discussing the manuscript.

