## Supplementary figures and images for "Microglial phagocytosis dysfunction is related to local neuronal activity in a genetic model of epilepsy"

### Supplementary Figure 1

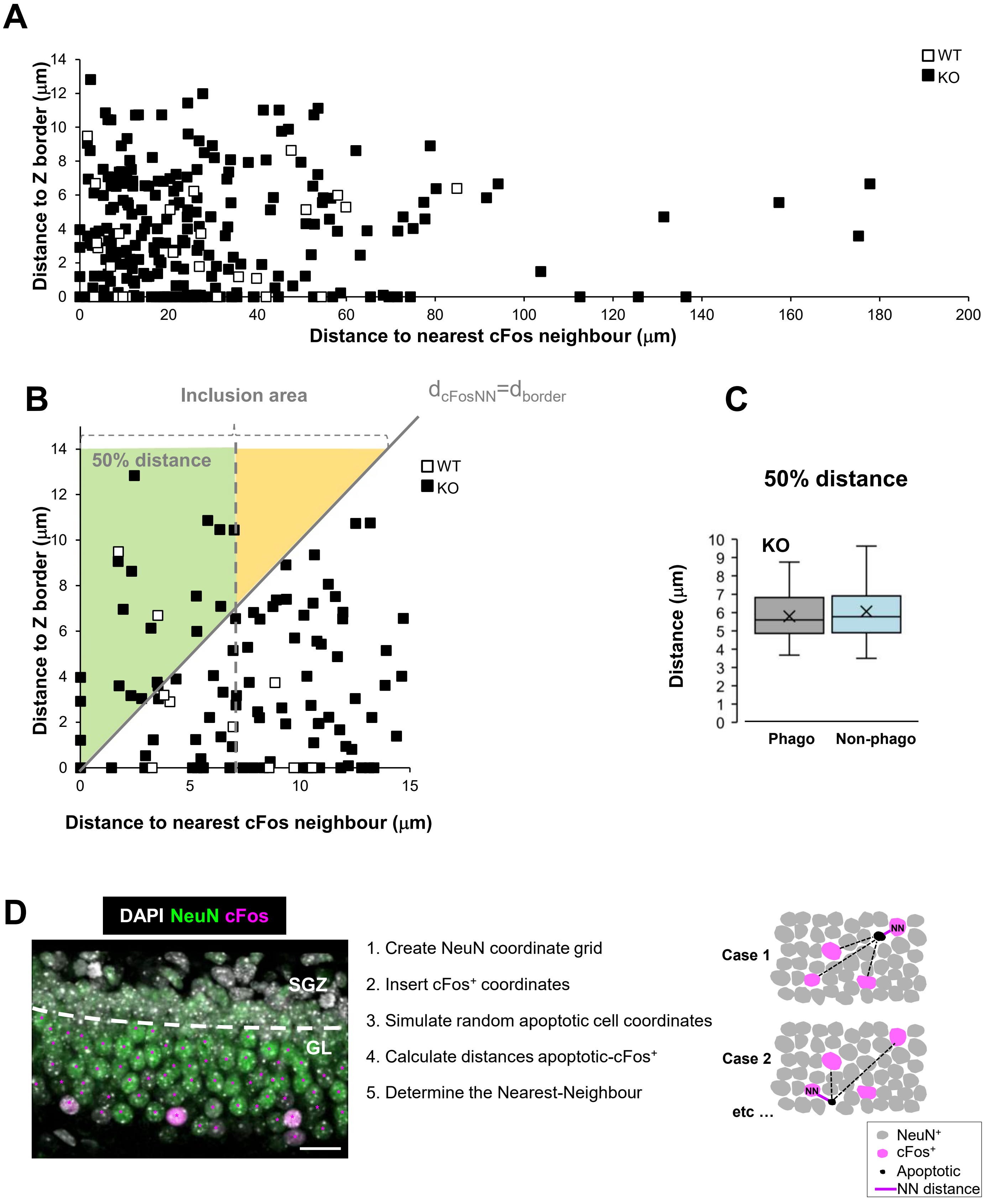
